# Genetic data and meteorological conditions: unravelling the windborne transmission of H5N1 high-pathogenicity avian influenza between commercial poultry outbreaks

**DOI:** 10.1101/2025.02.12.637829

**Authors:** Alexander Nagy, Lenka Černíková, Kamil Sedlák

## Abstract

Understanding the transmission routes of high-pathogenicity avian influenza (HPAI) is crucial for developing effective control measures to prevent its spread. In this context, windborne transmission, the idea that the virus can travel through the air over considerable distances, is a contentious concept and, documented cases are rare. Here, though, we provide genetic evidence supporting the feasibility of windborne transmission. During the 2023-24 HPAI season, molecular surveillance identified identical H5N1 strains among a cluster of unrelated commercial farms about 8 km apart in the Czech Republic. The episode started with the abrupt mortality of fattening ducks on one farm and was followed by disease outbreaks at two nearby high-biosecurity chicken farms. Using genetic, epizootiological, meteorological and geographical data, we reconstructed a mosaic of events strongly suggesting wind was the mechanism of infection transmission between poultry in at least two independent cases. By aligning the genetic and meteorological data with critical outbreak events, we determined the most likely time window during which the transmission occurred and inferred the sequence of infected houses at the recipient sites. Our results suggest that the contaminated plume emitted from the infected fattening duck farm was the critical medium of HPAI transmission, rather than the dust generated during depopulation. Furthermore, they also strongly implicate the role of confined mechanically-ventilated buildings with high population densities in facilitating windborne transmission and propagating virus concentrations below the minimum infectious dose at the recipient sites. These findings underscore the importance of considering windborne spread in future outbreak mitigation strategies.

## Introduction

Poultry farms are important emitters of air pollutants into the environment [1–3]. The airborne particles they emit vary in origin, size and shape, but all consist of a complex mix of gases, liquid droplets, and organic and inorganic matter [4]. Larger particles, visible to the naked eye, settle on surfaces as dust. The finest particles and liquid droplets, referred to as particulate matter, levitate in the air for extended periods and include size fractions that are easily inhalable and respirable.

During an outbreak of HPAI, viral particles can be regularly detected in the air and dust inside poultry houses or markets [5–11] and although airborne transmission is not considered the primary mode of infection in poultry [12, 13], contaminated air can contribute to the spread of the disease [14–16]. The question is to what extent HPAI viral particles can be carried by air currents?

Wind has long been considered a plausible factor in the long-distance transmission of the avian influenza A virus (IAV) [17]. However, field reports on this mode of spread are rare and air-sampling studies conducted around poultry or pig houses have not supported long-distance airborne IAV transmission. Such studies have reliably detected IAV only up to ∼150 meters away from infected farms, but with low viral loads and positivity rates [5, 6, 8, 9, 11, 18], and even fewer isolated viruses [5, 6, 18]. This suggests that virus load in the air is inversely proportional to the distance covered [19, 20]. Furthermore, the aerosol is a dynamic system where the interactions of environmental stressors, such as sunlight and ultraviolet radiation, temperature, relative humidity, and evaporation, with dispersed chemical and organic components, may rapidly inactivate IAV particles [21]. Consequently, with increasing distance, the contaminated plume emitted from a farm with an ongoing IAV outbreak is progressively diluted, and the proportion of infectious virions decreases. This mean that wind has been largely overlooked as a factor in IAV transmission and, thus, its influence perhaps underestimated during outbreak investigations.

Despite this, some reports have proposed that windborne spread plays a significant role in IAV transmission over longer distances under suitable weather conditions. In studies of the severe H7N7 HPAI outbreak in the Netherlands in 2003, one study claimed that wind spread accounted for 18% [20] and another 24% [22] of transmission events up to 25 km. Similarly, during the 2007 equine H3N8 influenza outbreak in Australia, 81% of infections within a cluster of 437 horse farms were attributed to windborne spread over a distance of 1-2 km [23]. Around the same time, the serological screening of turkeys in Minnesota in 2007-08 revealed that turkey premises within a 1.9 km radius of swine farms were most likely to test seropositive for H3N2 and H1N1 IAVs, thereby suggesting windborne transmission [24]. And during the 2014-15 multistate H5N2 HPAI outbreak in the USA, it was estimated that up to 39% of farms in Iowa alone could have experienced windborne infection within a radius of 8.5 km [25]. Overall, though, the discrepancy between air-sampling studies and transmission data raises questions about windborne IAV transmission. Over what distances can avian IAV be transmitted by the wind? Can air with even minute concentrations of avian IAV induce infection in a poultry flock?

Genetic data may prove crucial to answering such questions, but studies of windborne IAV transmission incorporating such data remain sparse. Batalie et al. [26] utilized partial genomic data to trace the transmission pathways of H7N7 HPAI during the 2003 Netherlands outbreak. For the same outbreak, Ypma et al. [20] used statistical correlations between the inferred routes of spread and weather conditions as genetic evidence of windborne transmission. Overall, though, there is a significant lack of genetic data mapped to the prevailing weather conditions for such outbreaks.

In this study, we present empirical genetic evidence supporting the windborne transmission of the H5N1 HPAI virus over a distance of 8 km. Molecular surveillance of HPAI outbreaks in the Czech Republic during the 2023-24 season identifies identical H5N1 strains in a cluster of unrelated commercial farms. This evidence, combined with a detailed epizootiological investigation and strong correlation with weather conditions, confirms that wind can transport infectious virus particles over substantial distances and, thereby, facilitate the spread of HPAI between poultry farms. These findings underscore the importance of considering windborne transmission as a significant factor in the management and control of HPAI outbreaks.

## Materials and Methods

### Whole-genome sequencing

Pooled organ suspensions, taken from each animal and homogenized in RNA Later solution (Invitrogen), were collected for next-generation sequencing. Total nucleic acids were extracted from 200 µl supernatants by MagNA Pure 24 (Total NA Isolation Kit), MagNA Pure 96 (DNA and Viral NA Small Volume Kit) (both from Roche), or MaxWell RSC (Maxwell® RSC Whole Blood DNA Kit, Promega) and eluted into 50 µl (MagNA Pure 24/96) and 60 µl (MaxWell RSC), respectively. The H5N1 genome was amplified in a two-step RT-PCR protocol. Reverse transcription was conducted using SuperScript III or SuperScript IV RT kits (Thermo Fisher Scientific) with a universal-tagged forward primer TTTCTGTTGGTGCTGATATTGC**AGCRAAAGCAGG** and utilizing a quarter of the recommended enzyme amount. Amplification was carried out with Q5 HiFi DNA polymerase (New England Biolabs; primers available on request) in a final volume of 25 µl (20 µl of master mix and 5 µl of RT reaction). The sequencing libraries were purified by SPRIselect magnetic beads (Beckman-Coulter) and quantified by QIAxpert (Qiagen). End preparation, native barcoding and sequencing adapter ligation were performed by Native Barcoding Kit (SQK-LSK-114.96, Oxford Nanopore Technologies (ONT)), according to the manufacturer’s instructions. Sequencing runs were carried out on the ONT MinION Mk1B using R10.4.1 flow cells and operated via MinKNOW software (v2.3.04.3 or higher, ONT). Basecalling was conducted with Dorado software (v0.5.0 or higher, ONT) using the super accurate basecalling model (dna_r10.4.1_e8.2_400bps_sup@v4.3.0 or higher model). The run was monitored in real time by the RAMPART (Read Assignment, Mapping and Phylogenetic Analysis in Real Time) module of the ARTIC bioinformatic pipeline [27] set to the concatenated H5N1 genome as a reference. Demultiplexing and adapter and barcode trimming were performed by Porechop. Consensus sequences were generated using various tools: Bcftools mpileup [28], NGSpeciesID [29], Decona [30], Amplicon sorter [31], and ViralConsensus [32]. The resulting consensus sequences were aligned using MAFFT (Multiple Alignment using Fast Fourier Transform) [33] and manually curated against the corresponding SAM files to obtain a final consensus genome. Finally, the obtained genomes were annotated using the Influenza Virus Sequence Annotation Tool [34]. All H5N1 genomes were submitted to the GISIAD EpiFlu [35] database with the accession codes listed in Table 1.

**Table 1.** Chronological list of H5N1 strains detected in the affected farms.

| Strain ID | Collection Date | Farm/House | Name | Accession no. |
| --- | --- | --- | --- | --- |
| 2268/24 | 2024-02-04 | B | A/domestic_duck/Czech_Republic/2268-1orig/2024 | EPI_ISL_18928997 |
|  |  |  | A/domestic_duck/Czech_Republic/2268-1/2024 | EPI_ISL_19338428 |
|  |  |  | A/domestic_duck/Czech_Republic/2268-2orig/2024 | EPI_ISL_18928998 |
|  |  |  | A/domestic_duck/Czech_Republic/2268-2/2024 | EPI_ISL_19338429 |
|  |  |  | A/domestic_duck/Czech_Republic/2268-3/2024 | EPI_ISL_19338430 |
|  |  |  | A/domestic_duck/Czech_Republic/2268-4/2024 | EPI_ISL_19338431 |
|  |  |  | A/domestic_duck/Czech_Republic/2268-5/2024 | EPI_ISL_18928999 |
|  |  |  | A/domestic_duck/Czech_Republic/2268-7/2024 | EPI_ISL_18929000 |
|  |  |  | A/domestic_duck/Czech_Republic/2268-9/2024 | EPI_ISL_19338432 |
| 2894/24 | 2024-02-12 | C1A | A/chicken/Czech_Republic/2894-1/2024 | EPI_ISL_19338427 |
|  |  |  | A/chicken/Czech_Republic/2894-4/2024 | EPI_ISL_18946721 |
|  |  |  | A/chicken/Czech_Republic/2894-5/2024 | EPI_ISL_18946722 |
| 2895/24 | 2024-02-12 | C2B | A/chicken/Czech_Republic/2895-1/2024 | EPI_ISL_18946713 |
|  |  |  | A/chicken/Czech_Republic/2895-2/2024 | EPI_ISL_18946712 |
|  |  |  | A/chicken/Czech_Republic/2895-3/2024 | EPI_ISL_18946716 |
|  |  |  | A/chicken/Czech_Republic/2895-4/2024 | EPI_ISL_18946715 |
|  |  |  | A/chicken/Czech_Republic/2895-5/2024 | EPI_ISL_18946714 |
| 2896/24 | 2024-02-13 | Backyard | A/chicken/Czech_Republic/2896-1orig/2024 | EPI_ISL_18946728 |
|  |  |  | A/chicken/Czech_Republic/2896-2orig/2024 | EPI_ISL_18946730 |
|  |  |  | A/chicken/Czech_Republic/2896-2/2024 | EPI_ISL_18946732 |
|  |  |  | A/chicken/Czech_Republic/2896-3/2024 | EPI_ISL_18946729 |
|  |  |  | A/chicken/Czech_Republic/2896-4/2024 | EPI_ISL_18946731 |
|  |  |  | A/chicken/Czech_Republic/2896-5/2024 | EPI_ISL_18946727 |
| 3299/24 | 2024-02-19 | C1B | A/chicken/Czech_Republic/3299-1/2024 | EPI_ISL_19000406 |
| 3529/24 | 2024-02-22 | C1B | A/chicken/Czech_Republic/3529/2024 | EPI_ISL_19044783 |
| 3627/24 | 2024-02-23 | C1B | A/chicken/Czech_Republic/3627-3/2024 | EPI_ISL_19000408 |
| 3549/24 | 2024-02-26 | C1B | A/chicken/Czech_Republic/3549-1/2024 | EPI_ISL_19000411 |
|  |  |  | A/chicken/Czech_Republic/3549-3/2024 | EPI_ISL_19338424 |
|  |  |  | A/chicken/Czech_Republic/3549-4/2024 | EPI_ISL_19044780 |
|  |  |  | A/chicken/Czech_Republic/3549-7/2024 | EPI_ISL_19338422 |
|  |  |  | A/chicken/Czech_Republic/3549-8/2024 | EPI_ISL_19338426 |
|  |  |  | A/chicken/Czech_Republic/3549-9/2024 | EPI_ISL_19044781 |
|  |  |  | A/chicken/Czech_Republic/3549-10/2024 | EPI_ISL_19044782 |
|  |  |  | A/chicken/Czech_Republic/3549-11/2024 | EPI_ISL_19000412 |
|  |  |  | A/chicken/Czech_Republic/3549-12/2024 | EPI_ISL_19338425 |
|  |  |  | A/chicken/Czech_Republic/3549-13/2024 | EPI_ISL_19338423 |
|  |  |  | A/chicken/Czech_Republic/3549-14/2024 | EPI_ISL_19000413 |
|  |  |  | A/chicken/Czech_Republic/3549-15/2024 | EPI_ISL_19000414 |

### Sequence and phylogenetic analysis

The concatenated H5N1 genomic sequences were aligned using MAFFT, with alignment trimming and format conversion (Phylip full names and padded) by AliView [36]. A maximum likelihood (ML) tree (IQ-TREE multicore version 2.2.0-beta for Linux 64-bit [37]; best fit moder GTR+F+G4 according to the Bayesian information criterion, 1,000 replicates) was calculated from the concatenated genomic segments. Segment concatenation was performed using the Union programme from EMBOSS [38] in the following order: PB2, PB1, PA, H5, NP, N1, MP and NS. A median-joining network was calculated using PopART (Population Analysis with Reticulate Trees) [39, 40] software with the ε set to zero.

### Meteorological data

Meteorological data (wind direction and speed, average temperature, and relative humidity), were obtained from three Czech Hydrometeorological Institute monitoring stations with the following IDs: B7TREB01, B2VMEZ01, and B2SEDC01.

## Results

### HPAI outbreak cluster of interest

On 4 February 2024, a commercial farm with ∼50,000 fattening ducks (hereafter referred to as B) experienced a sudden increase in mortality affecting ∼800 ducks in two of its eight houses. Within two days, the mortality rate increased to ∼5,000 deaths. The ducks were kept in identical houses on a littered floor with natural ventilation and a basic level of biosecurity. The farm was in close proximity to a ∼15 ha lake [41] accessed by wild birds. To control the HPAI outbreak, the entire flock was depopulated between 7 and 9 February, and a 3 km protection zone and 10 km surveillance zone established.

One week later, on 12 February 2024, H5N1 HPAI was detected in two chicken farms (hereafter referred to as C1 and C2) owned by another company, C. These farms were situated within the surveillance zone and, together with B, formed three vertices of an imaginary triangle (Fig 1). Company C is internationally recognised for its unique accredited breeding program, which supplies a variety of chicken hybrids with different plumage and egg colours to customers worldwide. Utilizing state-of-the-art breeding technology, this activity is carried out to the highest biosecurity standards.

**Fig 1.**
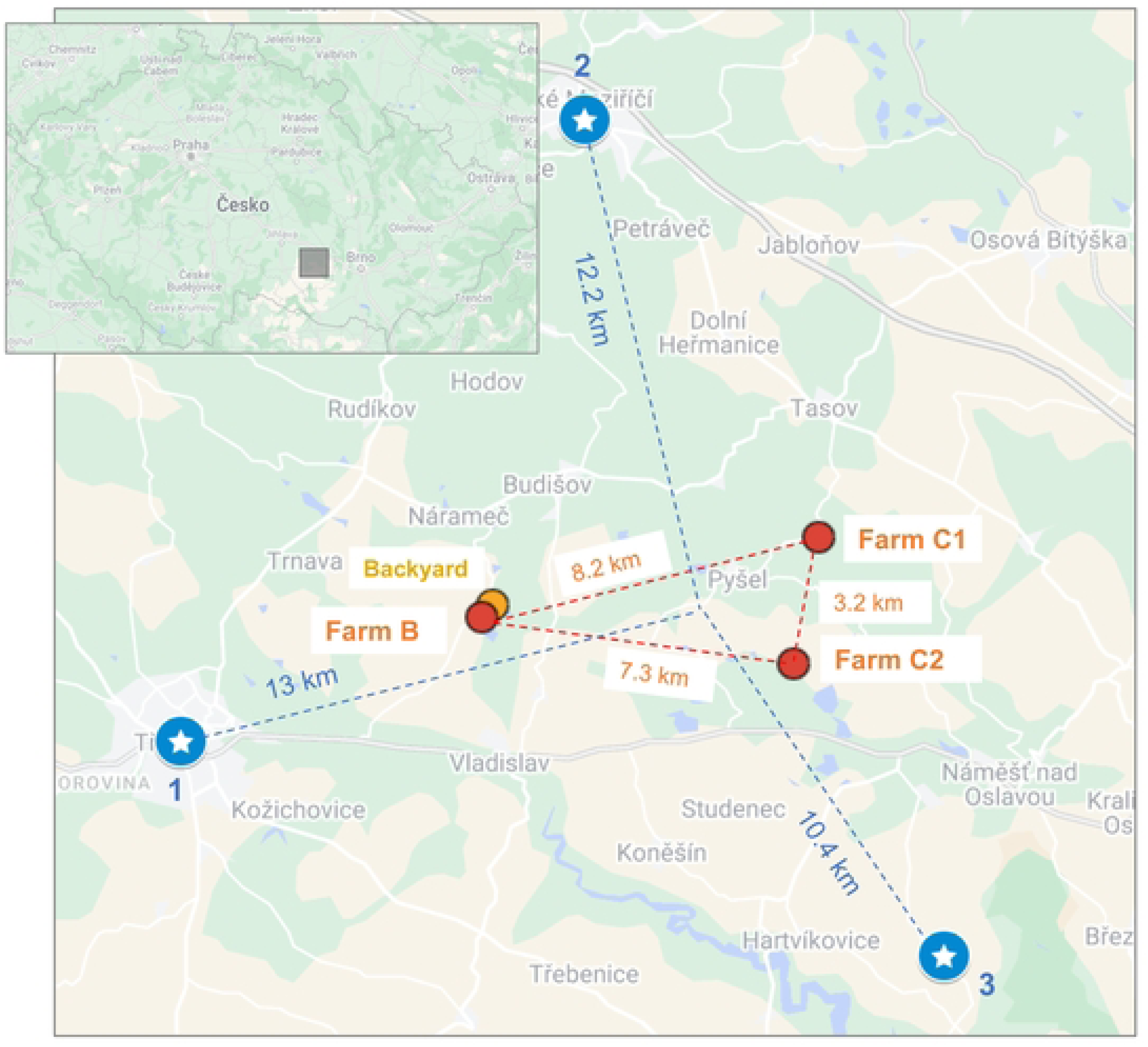
Map of outbreak locations. Commercial poultry farms are marked by red dots and backyard poultry farms by orange dots. Czech Hydrometeorological Institute monitoring stations are indicated by blue dots. An interactive version of the map is available online at: https://www.google.com/maps/d/edit?mid=1rlhK5gts6bwoNhAsamBt54wsvr8iBvc&ll=49.24410201219534%2C16.024409179698264&z=13.

Both C1 and C2 consisted of two houses, with birds kept in indoor cages without litter. C1 housed ∼24,500 birds: C1A ∼11,500 laying hens and C1B ∼13,000 birds of parent and grandparent breeds. C2 housed ∼45,000 birds: C2A ∼5,500 birds and C2B ∼39,500 birds intended for local sale. Both farms used treated water from their own wells and feed purchased from various companies. All houses were equipped with a negative pressure tunnel ventilation system (Fig 2). As the farms were within the surveillance zone, a Viusid Vet nutritional supplement (Catalysis, Spain) was administered to boost chicken immunity from 7 February 2024.

**Fig 2.**
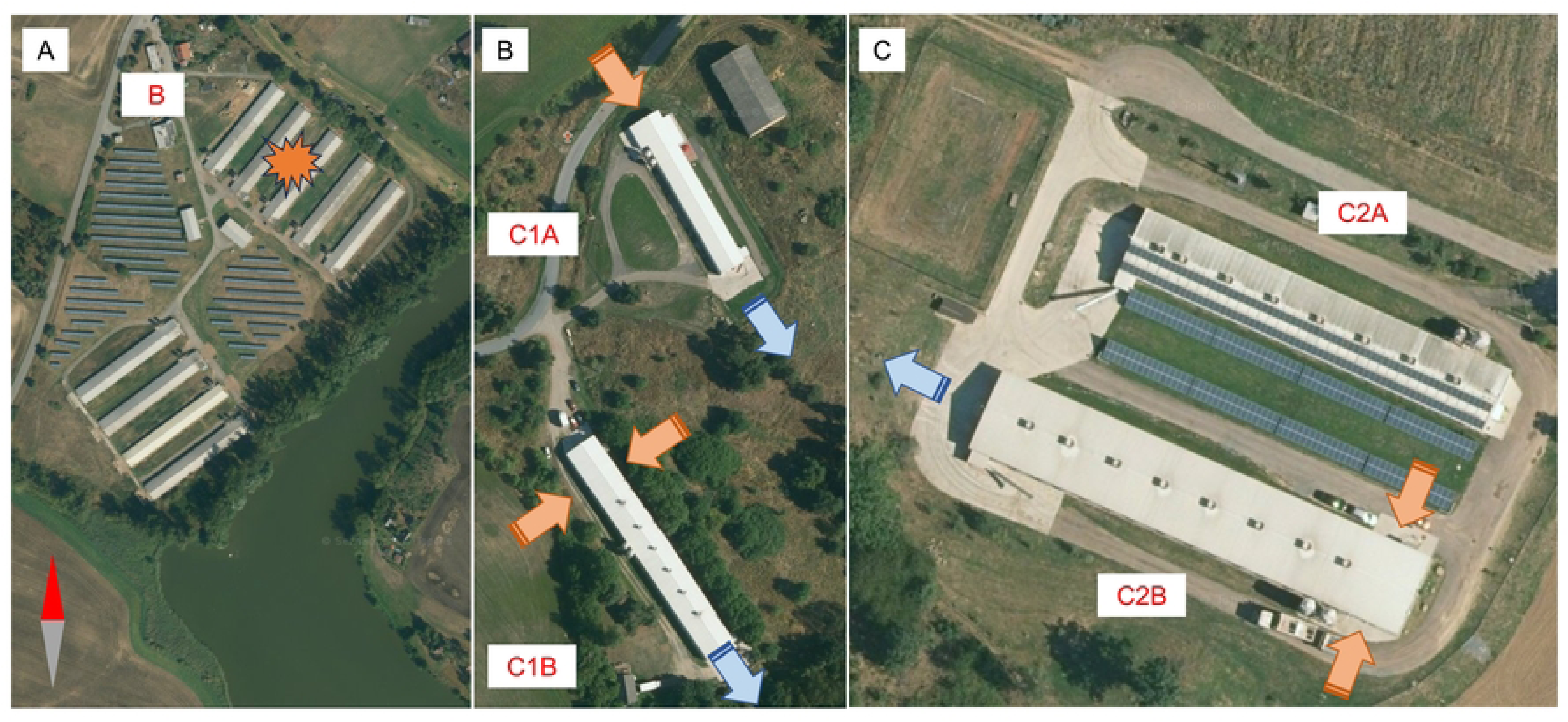
Infected Premises. Layout of affected houses on farms B (A), C1 (B), and C2 (C). On C1 and C2, tunnel ventilation system airflow is indicated with orange arrows for inflow and blue arrows for outflow.

H5N1 HPAI first manifested itself in C1A and C2B with an observed slight increase in mortality. For at least a week before HPAI was confirmed, both C1 and C2 experienced a gradual, though insignificant, decrease in water and feed consumption, with this decline initially attributed to the administration of Viusid. It is noteworthy that in the affected houses, the infection and subsequent mortality started in the areas closest to the air inlets. The situation in C1 was particularly serious as C1B contained ∼2,000 birds representing a crucial genetic reserve for the breeding program. To protect this indigenous gene pool, C1A was immediately depopulated. However, on 19 February, the virus was also detected in C1B. In an effort to limit the culling of this valuable population the progression of the infection was monitored (Fig 3). Ultimately, though, this house also had to be depopulated. C2A remained unaffected throughout.

**Fig 3.**
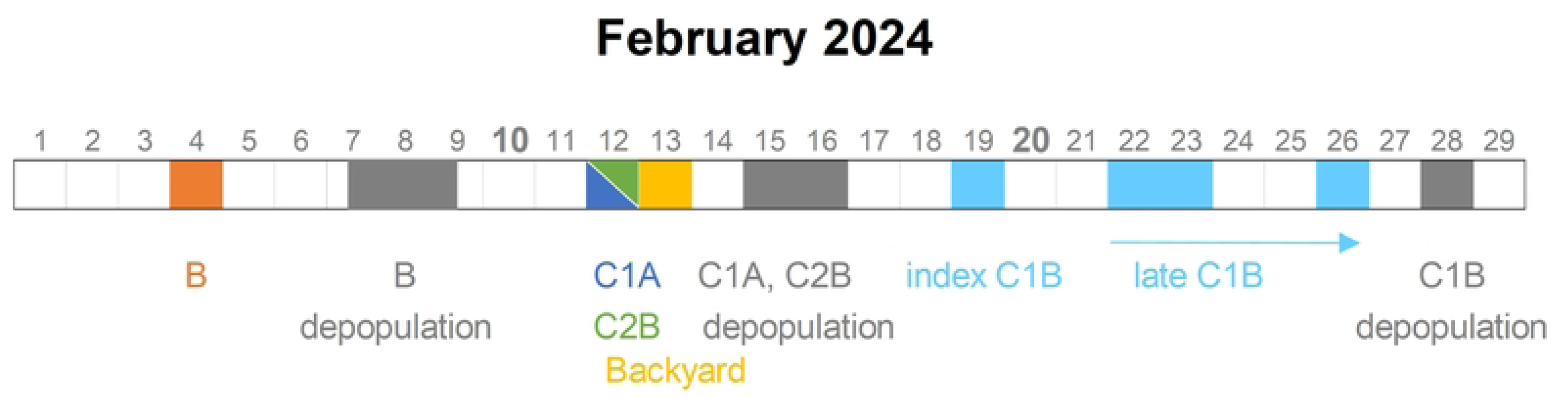
Time scheme of events. Timeline of key dates related to the outbreak, including outbreak identification, depopulation of affected premises, and specific sampling events carried out at the locations.

### Molecular epizootiology

To determine the relationships between the affected farms, whole-genome sequencing and subsequent phylogenetic analysis were performed. Total of 38 H5N1 genomes were obtained (Table 1), including nine genomes from B, three from C1A and five from C2B, each collected during a single sampling occasion. Another 15 were obtained from C1B during the course of four consecutive sampling events. An additional six genomes were obtained from the samples collected from the outbreak among backyard poultry near B.

Phylogenetic analysis showed that all H5N1 strains belonged to genotype DI [42] and formed a common sub-clade (Fig 4A). Interestingly, the H5N1 viruses from B and C1A and the index strain from C1B did not show farm-specific clustering but were placed in statistically unsupported branches. In particular, three H5N1 strains from B were 100% identical at the nucleotide level to strains from C1A and the index C1B. In contrast, H5N1 strains from C2B and the later strains from C1B showed clear farm-specific clustering. Finally, H5N1 viruses from backyard poultry identified nine days apart near B (Figs 1, 3), showed a close relationship to the B, C1A and index C1B strains.

**Fig 4.**
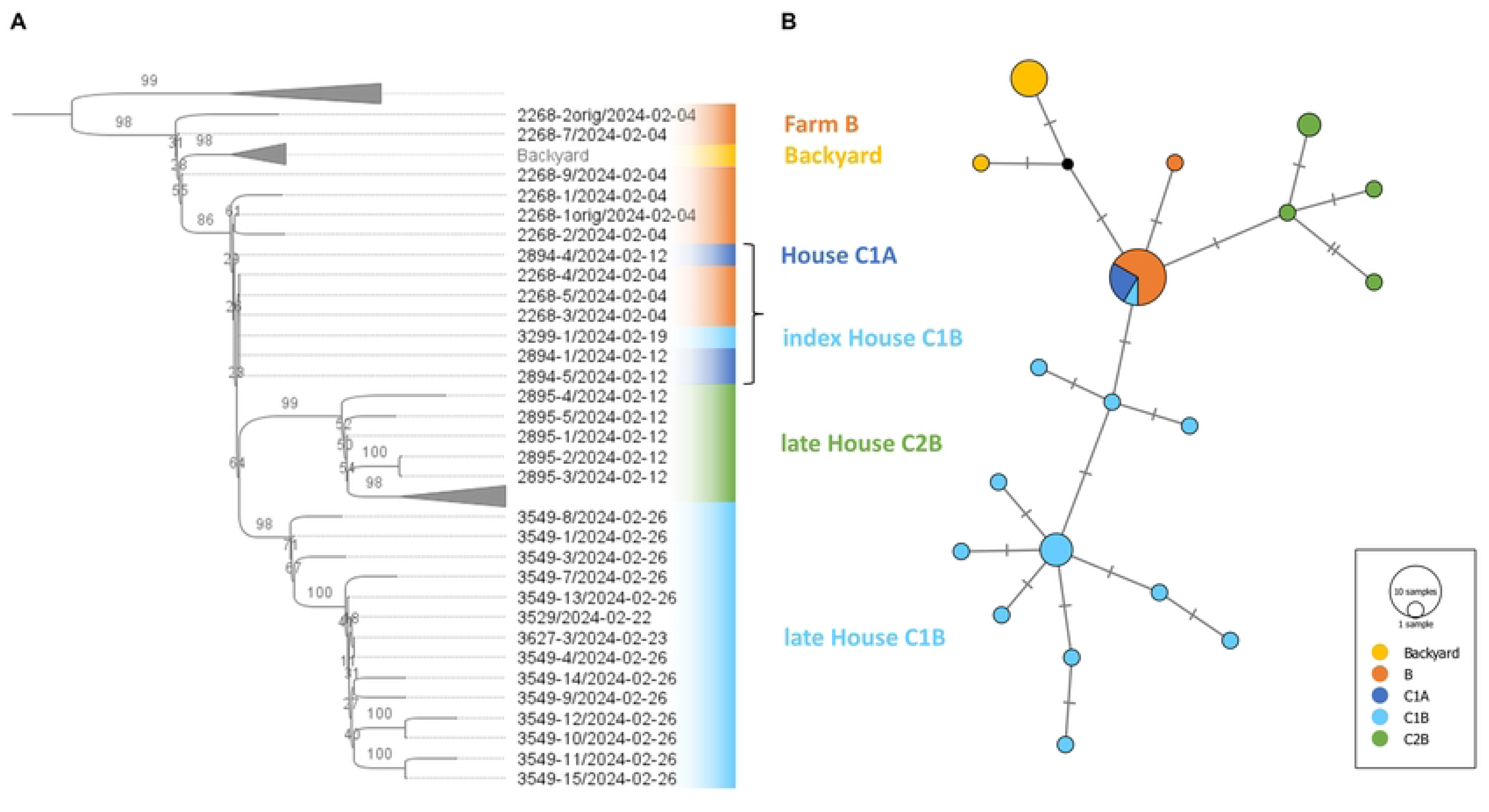
Phylogenetic analysis. Maximum likelihood tree (A) constructed using concatenated genomic data. Each branch is annotated with bootstrap values (derived from 1000 replicates) expressed as percentages. Genomes with 100% sequence identity are indicated by a square bracket. Median-joining network (B) of the same dataset. The sizes of the vertices are proportional to the number of genomes included. Nucleotide differences are shown as hatch marks on the edges.

To further elucidate the relationship between B, C1 and C2, a median-joining network was calculated. This revealed that all but one of the H5N1 strains from B and the early H5N1 variants from C1 were grouped into a single central node (Fig 4B). From this node, three main branches emerged: the first two represented C1B- and C2B-specific clusters, and the third grouped H5N1 strains from backyard poultry. These results are consistent with those obtained from the genomic tree. This genetic overlap between B and C suggests a common origin, with B likely being the primary source from which the disease spread to C1 and C2.

### Tracing the potential infection routes

Companies B and C were only about 8 km apart and active in different market segments. Interviews with the veterinary inspector and general manager of C revealed the complete absence of any interaction between the companies, even through third parties involved in feed replenishment, waste disposal or the transport of carcasses to rendering plants. All farms used their own well water supplies. In addition, the employees were not allowed to keep their own poultry. Therefore, in view of the biosecurity measures in place, the possibility of human-associated secondary spread from B to C1 and C2, or between C1 and C2 can be excluded.

Because C1 and C2 were located in an area without significant water bodies, H5N1 transmission from wild birds also appears highly unlikely. The houses were enclosed in a well-maintained clean environment with reinforced fencing that effectively prevented the entry of rodents and small carnivores known to be susceptible to the virus [43–46]. Furthermore, the combination of the outbreak season and tunnel ventilation system ruled out the involvement of flies or other insects in infection transmission.

Thus, the genetic data and field investigations suggest an abiotic route of transmission from B to C1 and C2.

### Local landscape and meteorological conditions

All farms were situated at similar altitudes: B at 438 m, C1 at 448 m and C2 at 408 m (Fig 5A). The landscape between the farms consisted mainly of open countryside, fields, meadows and sparse tree cover with the highest local hill being 510 m.

**Fig 5.**
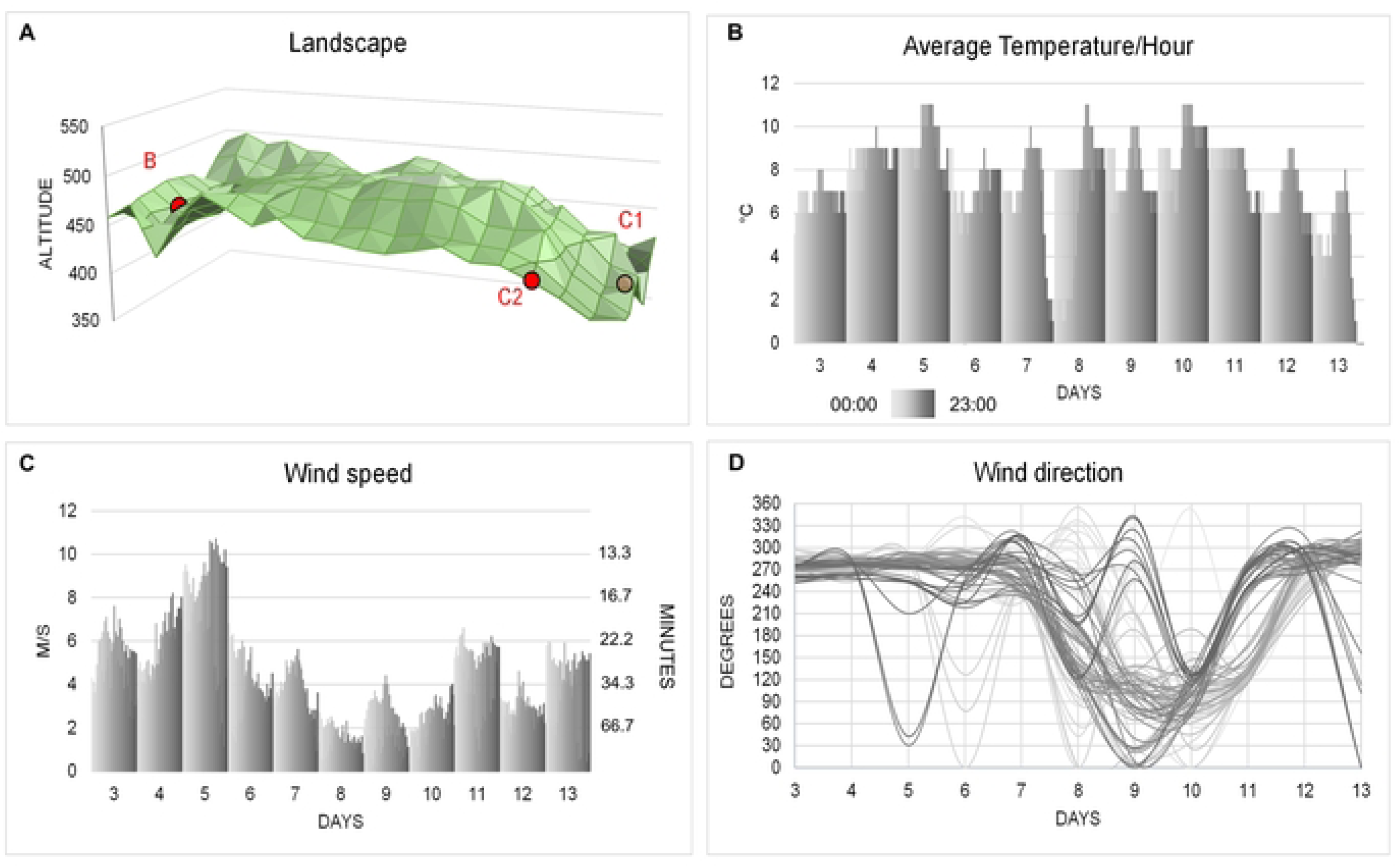
Landscape and meteorological conditions. Landscape (A) characterises the ruggedness of the terrain between the donor and recipient farms according to altitude in metres above sea level. Average temperature (B), wind speed (C) and wind direction (D) all recorded at hourly intervals at three Czech Hydrometeorological Institute stations. Temperature was recorded at two metres above ground, and wind speed / direction at ten metres. Time data from 0:00 to 23:00 h are shaded in grey. The secondary y-axis (C) indicates the time to reach recipient farm C1 at a given wind speed.

The prevailing weather conditions included extensive cloud cover, minimal precipitation and intermittent sunshine. Humidity levels averaged from 77 to 81%. Apart from the night of 7 to 8 February, temperatures did not fall below 6 °C, peaking at 8-11 °C on 4 and 5 February. Wind conditions (Fig 5B-D) were remarkable between 4 and 7 February, with continuous wind from the west or southwest (250-300 degrees). Generally, wind speeds were around 4 m/s (∼14 km/h), but there were short periods of turbulence during which they increased to over 6 m/s (∼22 km/h) on 4 February and reached 8-10 m/s (∼29-34 km/h) on 5 February. Thereafter, the wind velocity decreased and became erratic, sometimes coming from the east and southeast on 8 and 9 February. Significantly, the weather conditions correlated with the transmission route inferred from the genetic data, with the optimal period for infection spread being from noon on 4 February to midnight on 5 February.

### Synthesis of key findings suggests windborne transmission

The H5N1 HPAI outbreak started at B on 4 February 2024. Although the exact source of the infection remains unidentified, it most likely originated from mallards at a nearby pond. While this hypothesis lacks genetic confirmation, both the sudden onset of the disease and the mortality rate in the commercial ducks suggest a high infectious dose delivered in a short period, likely by wild waterfowl. The fact that a nearby backyard flock was subsequently (around nine days later) infected with genetically similar H5N1 strains further indicates that wild birds sustained virus circulation in the area.

Alignment of the events timeline with wind direction and speed suggest a six-day window (from outbreak identification on 4 February to depopulation completion on 9 February) during which the infection was spread from B to C1 and C2. For this window, the theory of windborne spread, which began with an order of magnitude increase in the number of infected birds at B, is supported by three key observations: i) slow disease progression with a relatively long incubation period characterised by a decrease in water and feed consumption before the clinical symptoms became overt, ii) the location of the affected birds in sections closest to the air inlets [25], and iii) the genetic identity between the H5N1 strains in the donor and recipient farms. Moreover, all possible alternative routes of infection during this period were excluded by our field investigation.

Thus, the C1A and index C1B viruses were most likely seed strains carried by wind from B. These seed strains then evolved locally, forming a cluster of farm-specific variants [26] detected in C2B and later C1B. While the critical early phase of infection was not captured for C2B, the timeline of events and H5N1 sequence diversity suggest infection in the following order: C2B, C1A and C1B. However, it remains unclear whether C1B was infected separately or through ventilation output from C1A (Fig 2).

## Discussion

Understanding the routes of IAV transmission is crucial for developing strategies to control its spread and prevent both inter-species and cross-species infections. While the windborne spread of IAV is considered a possible route for infecting poultry, pigs and even horses, the unpredictable nature of infection outbreaks, the specific weather conditions required and the lack of samples from crucial transmission time points make it difficult to confirm this mode of spread. Consequently, our ability to infer windborne spread in the field has largely been limited to correlation or simulation studies performed retrospectively [22–25, 47, 48]. There is a lack of robust genetic data to support these inferences.

In the current study, we have supported previous findings [20, 26] by providing direct genetic evidence for the windborne transmission of H5N1 HPAI from whole-genome sequence data. Our analysis indicates that the high H5N1 genomic identity between the unrelated poultry farms can only be attributed to an abiotic route. The combination of landscape, meteorological, clinical and epizootiological factors points to wind-mediated spread as the only plausible explanation.

The windborne spread of H5N1 HPAI in the cluster of interest raises significant questions about the infection dynamics of housed poultry, particularly with respect to the role of tunnel ventilation systems in facilitating such infection. Tunnel ventilation systems in poultry houses act like high-volume air samplers; powered by exhaust fans, they create a negative pressure that pulls in ambient air [49]. The maximum ventilation capacity, estimated in cubic meters per hour per kilogram of body weight, usually ranges between 4.5 and 7 m^3^ [50]. For a poultry house with 10,000 chickens, each weighing 1 kg, this results in the influx of a colossal volume of 45,000 to 70,000 m³ of ambient air per hour, which is equivalent to a cube with an edge ranging from 34 to 41 meters. This enables a significant amount of particulate matter to enter the chicken house. During stormy periods, the influx can be even greater [51]. Consequently, inside such a poultry house, the dense population of birds is permanently exposed to large volumes of ambient air. Indeed, the prolonged exposure of immunologically naive chickens to contaminated air, even at virus concentrations below the minimal infectious dose (MID), has been shown to may increase the risk of airborne infection [25, 48].

During the infection period, the ventilation rates in C1A, C1B and C2 were set to a staggering 154,111, 134,228 and 257,244 m³/h, respectively. The exposure of a dense population of chickens to such high air volumes might well lead to minute virus amounts being captured and subsequently enriched to levels sufficient to trigger illness. This is consistent with the observed slow disease progression. Thus, it is likely that the specific settings of the poultry houses at recipient site C facilitated the wind-mediated spread of HPAI. Conversely, this also explains why other backyard poultry in villages located between the donor and recipient farms, as well as the poultry in C2A, were unaffected; they lacked sufficient population density and/or the booster effect of tunnel ventilation. Furthermore, the population-dependent and booster effects also align with the deduced order of infected farms, indicating that C2B, with the largest bird population and highest ventilation rate, was infected first. Therefore, while further studies are needed, it does appear that wind can transmit infectious HPAI particles over considerable distances and that densely populated tunnel-ventilated poultry houses can promote the propagation of sub-MID HPAI virus concentrations.

However, detecting viral particles in such minute concentrations is challenging. Simply put, at greater distances, virus amounts are below the detection limit of existing air samplers, making reliable estimation extremely difficult, if not impossible, as discussed by Zhao et al. [25]. In addition, given the invisibility of the plume and the unpredictability of air currents, measurements have to be performed at the right time and place under weather conditions conducive to windborne spread. Accordingly, and consistent with Zhao et al. [25], negative results from air samples collected at greater distances may not accurately reflect an IAV-negative air environment. Thus, the failure to detect IAV particles away from infected farms [5, 6, 8, 9; 11] must not be taken as evidence of the infeasibility of windborne spread. Indeed, when sampling was correlated with careful estimation of wind direction, IAV particles were detected in air collected up to 1.5 and 2.1 km from affected farms, indicating true wind-mediated dispersal [18].

By aligning the timeline of events with wind velocity and a detailed knowledge of the clinical progression, we are able to infer with high confidence the time window during which the infection was transmitted by the wind from B. The period from outbreak identification to the end of depopulation (3‒9 February) leaves about seven days during which the virus could have been carried to C1 and C2. Taking into account the wind velocity, with an almost constant easterly flow between 4 and 7 February, this window can be further reduced to four days. However, if the observed decrease in water and feed intake is considered to be the first sign of infection [52], then infection occurred between 4 and 5 February. This period corresponds to the rapid increase in the number of infected birds on the donor farm, which must have acted as a massive source of aerosolised virus particles. Additionally, the highest wind speeds were recorded for a period of about 36 h (noon on 4 February to midnight on 5 February), enabling the virus to reach C1 and C2 within 13-22 min. Furthermore, correlating the observed genomic diversity with the sampling dates shows that these houses were infected independently, highlighting the underlying infectious dynamics. Overall, it seems that the recipient farms were infected relatively very early after the outbreak on the donor farm and before it was depopulated. These findings are in stark contrast to the conventional theory [53, 54] that dust generated during depopulation is the major risk factor for the secondary wind spread of HPAI. It appears that the smoother aerosols produced by an infected flock are much more likely carriers than the far heavier dust particles that settle quickly within short distances of an outbreak farm.

Windborne transmission is a controversial concept in relation to the spread of avian influenza. While some studies suggest that it can play a significant role, others consider the potential for airborne transmission low. The specific, often ambiguous, conditions required make long-distance spread seem counterintuitive. As a result, it is questionable whether the windborne route has been adequately considered during outbreak investigations, meaning that, compared with other transmission routes, it may well have been underestimated. However, when factors, such as suitable weather conditions, proximity to the source population and densely populated tunnel-ventilated houses, align, windborne transmission becomes a feasible mode of spread over kilometric distances, as demonstrated by at least two independent events observed in our study. It is therefore worth considering the integration of airborne control technologies [55] with existing on-farm biosecurity protocols to effectively limit the introduction and spread of pathogens via the windborne route. Continued research and improved surveillance are essential to fully understand the risks associated with such transmission. The knowledge gained could then be used to develop more effective preparedness strategies and control measures that would better mitigate the effects of future outbreaks.

## Acknowledgements

We wish to thank Kristína Oreničová, B.Sc., Michaela Štěrbová, B.Sc., Lada Hofmannová, DVM, PhD, Ondřej Horák, DVM, PhD, Roman Maleček and all our colleagues at the State Veterinary Institute Prague and Jihlava for their excellent technical support in conducting the HPAI survey. We would also like to thank Marie Nováková, DVM, from the Regional Veterinary Administration of the Vysočina Region, and Pavel Dobrovolný, Managing Director of company C, for providing valuable information from the field. In addition, we are grateful for the bioinformatics assistance of Petr Buřič, M.Eng.

